# Instantaneous Frequency: A New Functional Biomarker for Dynamic Brain Causal Networks

**DOI:** 10.1101/2024.12.17.628965

**Authors:** Haoteng Tang, Siyuan Dai, Lei Guo, Pengfei Gu, Guodong Liu, Alex D. Leow, Paul M. Thompson, Heng Huang, Liang Zhan, the Alzheimer’s Disease Neuroimaging Initiative

## Abstract

This study introduces instantaneous frequency (IF) analysis as a novel method for characterizing dynamic brain causal networks from fMRI blood-oxygen-level-dependent (BOLD) signals. Effective connectivity, estimated using dynamic causal modeling (DCM), is analyzed to derive IF sequences, with the average IF across brain regions serving as a potential biomarker for global network oscillatory behavior. Analysis of data from the Alzheimer’s Disease Neuroimaging Initiative (ADNI), Open Access Series of Imaging Studies (OASIS), and Human Connectome Project (HCP) demonstrates the method’s efficacy in distinguishing between clinical and demographic groups, such as cognitive decline stages, sex differences, and sleep quality levels. Statistical analyses reveal significant group differences in IF metrics, highlighting its potential as a sensitive indicator for early diagnosis and monitoring of neurodegenerative and cognitive conditions.

**Graphical Abstract:** 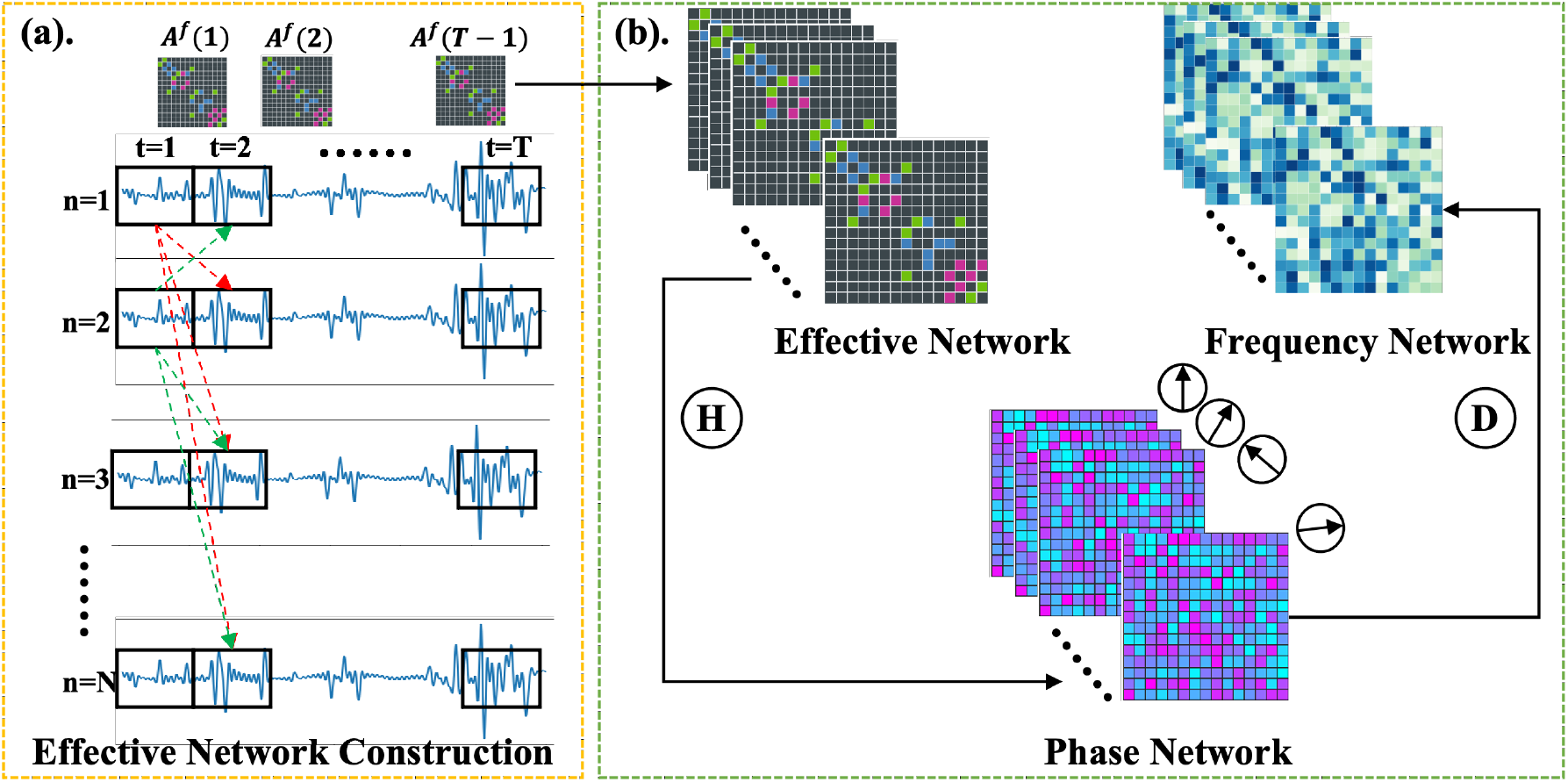

**Highlights:**

- The study introduces instantaneous frequency (IF) as a novel biomarker derived from dynamic brain effective connectivity, capturing temporal fluctuations in brain networks.
- The proposed IF biomarker effectively differentiates between various clinical stages, such as Mild Cognitive Impairment (MCI) and Alzheimer’s Disease (AD), and demographic factors, including sex and sleep quality.
- The robustness and clinical relevance of the IF biomarker are validated using three independent datasets: ADNI, OASIS, and HCP, demonstrating its potential in cognitive and neurological research.

## 1. Introduction

In computational neuroscience, brain biomarkers serve as essential quantifiable indicators of biological processes, aiding in detecting, characterizing, and monitoring brain pattern changes, progressing neurological conditions, and mechanisms underlying various clinical phenotypes. These biomarkers provide a window into complex brain dynamics, supporting precision in understanding disease development and individual variability across clinical states. Structural biomarkers, generally measuring stable brain anatomical features such as gray matter volume, cortical thickness, and white matter integrity, provide valuable insights into long-term and static neural integrity and structural abnormalities that accompany neurological conditions [1, 2]. However, structural metrics, while informative, often fall short in capturing the dynamic and activity-driven aspects of brain function associated with complex cognitive and pathological states. Temporal neural interactions and adaptive responses offer insights into how brain regions coordinate over time, revealing mechanisms and compensatory functions crucial for studying mental and neurological disorders [3, 4, 5, 6]. As a result, research has increasingly focused on functional biomarkers that capture dynamic changes in neural responses fluctuating with cognitive and pathological demands, and provide a nuanced view of real-time interactions among different brain regions. By capturing temporal relevance and causative changes, functional biomarkers facilitate early diagnosis, reveal disease heterogeneity, and enable new insights into disease progression and cognitive variability [7, 8].

Dynamic causality across brain regions is a critical component of brain functionality, encompassing the directional interactions between distinct brain regions and offering insights into how these interactions evolve and adapt over time [9, 7]. This causal fluctuation is integral to understanding not only the basic mechanisms of brain connectivity but also its perturbations among different neurodegenerative stages, such as Mild Cognitive Impairment (MCI) and Alzheimer’s Disease (AD) [10, 11] These fluctuations reveal essential differences in the adaptability, resilience, and stability of brain networks under both healthy and pathological conditions. Furthermore, dynamic causality is associated with various clinical phenotypes, including gender-related neural differences [12] and the impact of sleep quality on brain connectivity [13]. By examining these interactions, researchers can uncover critical markers of brain health and dysfunction. To investigate these dynamic causal interactions, this study employs Blood Oxygen Level-Dependent (BOLD) signals derived from functional magnetic resonance imaging (fMRI). BOLD signals provide a non-invasive method for measuring neural activity, allowing for the inference of effective connectivity through computational models such as the dynamic causal model (DCM) [14]. DCM is a well-established framework that estimates the time-dependent, directional influences that one brain region exerts on another, offering a comprehensive view of brain network dynamics [15].

In this study, we introduce a novel approach to derive functional biomarkers from temporal brain effective networks by analyzing the rate of change in causal influences across brain regions. We compute the instantaneous frequency (IF) of these effective connectomes, capturing time-dependent fluctuations in effective connectivity, and define the average instantaneous frequency across nodes as our biomarker. This measure reflects the oscillatory behavior of the entire network, offering insight into global brain dynamics. Our experimental results indicate that this biomarker exhibits significant differences across neurodegenerative stages, such as Alzheimer’s Disease (AD) and Mild Cognitive Impairment (MCI), as well as among other clinical phenotypes. To enhance the specificity of our findings, we conduct connectome-level analyses, pinpointing particular connectomes with significant clinical phenotype-related differences. This connectome-focused analysis allows us to identify targeted connections within the network that serve as precise markers of brain state changes underlying various clinical phenotypes and neuropsychiatric conditions.

The main contributions of this work are summarized as follows: (1) We introduce a novel biomarker, termed instantaneous frequency (IF), derived from brain effective networks to quantify and model functional fluctuations in the human brain. (2) We demonstrate that the proposed IF metric exhibits significant differences across various brain disorder stages and clinical phenotypes, providing a sensitive indicator for distinguishing these conditions. (3) Utilizing the proposed IF, we identify distinct brain connectomes associated with specific clinical and pathological groups, shedding light on the neural mechanisms underlying these variations.

## 2. Methods

### 2.1 Preliminaries

A brain effective network with *N* nodes is a weighted undirected graph *G* = {*V, E*} = (*A, X*), where 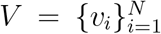 is the set of graph nodes representing brain regions, and *E* = {*e*_*i,j*_} is the directed edge set. *X* ∈ ℝ^*N* ×*c*^ is the node feature matrix where *x*_*i*_ ∈ ℝ^*c*^ is the *i*−th row of *X* representing the node feature of *v*_*i*_. *A* ∈ ℝ^*N* ×*N*^ is the adjacency matrix where *a*_*i,j*_ ∈ ℝ represents the weights of the edge between *v*_*i*_ and *v*_*j*_. As brain effective network is an undirected graph, *a*_*i,j*_ ≠ *a*_*j,i*_ ∈ ℝ. The sign of *a*_*i,j*_ indicates the direction of causal impact between *v*_*i*_ and *v*_*j*_, where *a*_*i,j*_ *>* 0 signifies the causal effect on *v*_*j*_ induced by *v*_*i*_, vice versa. Additionally, we denote the blood-oxygen-level-dependent (BOLD) signal obtained from fMRI as *B* ∈ ℝ^*N*×*b*^, where *b* is the length of the signal.

### 2.2 Construction of Dynamic Brain Effective Network

We construct brain effective networks by utilizing BOLD signals obtained from functional MRI with a dynamic causal modeling (DCM) framework [16, 17]. Each brain region is represented as an effective network node, while temporal changes in effective connectivity define the edges of the network. This temporal effective connectivity, represented by the dynamic adjacency matrix, *A*(*t*) can be modeled as:

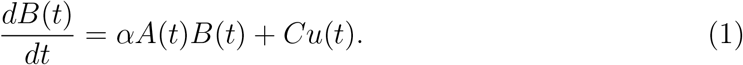

Here *Cu*(*t*) accounts for external neuronal inputs, *u*(*t*), that influence dynamic *A*(*t*)). For our purposes, *Cu*(*t*) = 0 since we focus on resting-state fMRI, eliminating external influences. The parameter *α* is a constant modulating neuronal lag among brain nodes. Consequently, we can derive the expression of *A*(*t*) as follow:

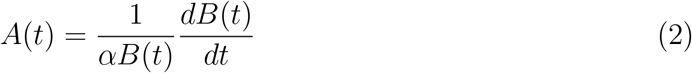

To discretize this continuous model, we define *A*(*t*)) in terms of BOLD signal change over discrete time intervals, resulting in:

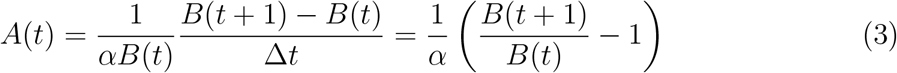

The effective connectivity between nodes 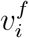 and 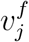 at time *t* is defined as follow:

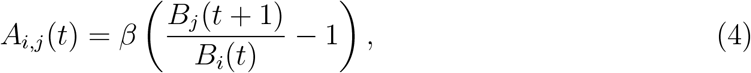

where 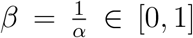 and *B*_*i*_ is the BOLD signal at node *v*_*i*_. This framework allows us to analyze effective connectivity dynamics in resting-state brain networks, providing insights into causal interactions across brain regions.

### 2.3 Instantaneous Frequency of Effective Connectomes

In the adjacency matrix, *A*(*t*) ∈ ℛ^*N* ×*N* ×*b*^, of the brain effective network, each element *A*_*i,j*_(*t*) ∈ ℛ^*b*^ reflects the time-varying casual impact between node *v*_*i*_ and *v*_*j*_. We first perform Hilbert transform to each *A*_*i,j*_(*t*) to generate the phase matrix, and then we compute the instantaneous frequency matrix by utilizing the phase matrix [18, 19, 20].

#### 2.3.1 Computation of Instantaneous Phase Matrix

Denote the instantaneous phase matrix as *P* ∈ (*t*) ℛ^*N*×*N*×*b*^, which provides the instantaneous phase of dynamic effective connectivity over time. The instantaneous phase signal for each dynamic effective connectivity can be computed as follow:

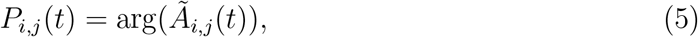

where Ã_*i,j*_ (*t*), representing the analytic signal of Hilbert transform *H*[·], can be computed as follow:

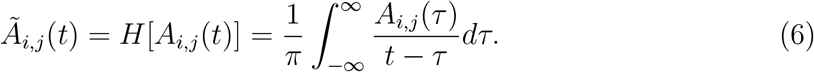

#### 2.3.2 Computation of Instantaneous Frequency Matrix

To obtain the instantaneous frequency matrix *F* (*t*), we compute the time derivative of each element in the phase matrix *P* (*t*):

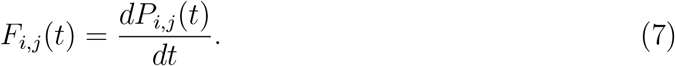

This instantaneous frequency represents the rate of change in causal influence between nodes over time. It captures the fluctuation speed and frequency of causal relationships, indicating the dynamic regulatory properties of these effective connections. High instantaneous frequency indicates rapid changes in causality, implying a high level of flexibility or adaptability in brain networks under certain conditions.

#### 2.3.3 Computation of Functional Biomarker

To compute the functional biomarker, Ω(*t*), of the global effective network, we average the node-level instantaneous frequency sequence:

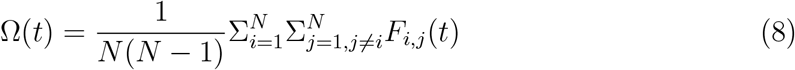

### 2.4 Statistical Method for Group Difference Analysis

Since the functional biomarkers Ω ∈ ℛ^*b*^ are time-series data, we use the Dynamic Time Warping (DTW) distance [21, 22, 23] to assess the similarity between each pair of sequences. For two distinct groups of biomarkers, we calculate intra-group DTW distances within each group and a cross-group DTW distance between them. To evaluate statistical significance, we apply two Mann-Whitney U tests [24, 25] comparing each intra-group distance against the cross-group distance. If both tests indicate significant differences, we consider the two groups of biomarkers to be significantly different.

## 3. Results and Discussions

### 3.1 Data Description and Preprocessing

This study employs three independent brain imaging datasets. The first dataset is from Alzheimer’s Disease Neuroimaging Initiative (ADNI) [26]. The ADNI was launched in 2003 as a public-private partnership, led by Principal Investigator Michael W. Weiner, MD. The original goal of ADNI was to test whether serial magnetic resonance imaging (MRI), positron emission tomography (PET), other biological markers, and clinical and neuropsychological assessment can be combined to measure the progression of mild cognitive impairment (MCI) and early Alzheimer’s disease (AD). The current goals include validating biomarkers for clinical trials, improving the generalizability of ADNI data by increasing diversity in the participant cohort, and to provide data concerning the diagnosis and progression of Alzheimer’s disease to the scientific community. For up-to-date information, see (http://adni.loni.usc.edu). Our ADNI data includes 154 normal control (NC) subjects (mean age = 77.46 ± 5.98, 59 males), 70 early mild cognitive impairment (EMCI) subjects (mean age = 73.75 ± 5.72, 38 males), and 50 late mild cognitive impairment (LMCI) subjects (mean age = 73.86 ± 6.20, 27 males). The second dataset is from the OASIS-3 project (Open Access Series of Imaging Studies-3), a longitudinal, multimodal resource encompassing neuroimaging, clinical, cognitive, and biomarker data focused on normal aging and Alzheimer’s disease. Our OASIS data includes 1051 NC subjects (mean age = 69.17 ± 8.90, 425 males), 209 MCI subjects (mean age = 75.44 ± 6.90, 122 males), and 66 Alzheimer’s Disease (AD) subjects (mean age = 74.29 ± v8.94, 41 males). The third dataset is from the Human Connectome Project (HCP)[27], a large-scale neuroscience initiative that maps the brain’s structural and functional connectivity in unprecedented detail. It serves as a comprehensive resource for understanding how different regions of the human brain are interconnected and how these connections support cognition, behavior, and neural function. Our HCP data includes 1206 young healthy subjects (mean age 28.19 ± 7.15, 603 women).

### 3.2 Significant Analysis of Functional Biomarker

#### 3.2.1 Experiment Description

In this section, we conduct a comprehensive analysis across multiple datasets to investigate if our proposed biomarker can serve as an effective indicator to distinguish various clinical and demographic groups. By investigating the biomarker across these various datasets and group comparisons, we aim to validate its robustness and relevance as a tool for understanding cognitive and clinical variability within and across different populations.

For the ADNI dataset, we analyze differences in the biomarker across three distinct subject groups: the Normal Control (NC) group, representing cognitively healthy individuals; the Early Mild Cognitive Impairment (EMCI) group, which includes participants with early-stage cognitive decline; and the Late Mild Cognitive Impairment (LMCI) group, representing individuals with more advanced cognitive impairment. By comparing these groups, we aim to evaluate the biomarker’s sensitivity to different stages of cognitive decline, capturing its potential responsiveness to progressive changes across the continuum of cognitive impairment.

In our analysis of the OASIS dataset, we similarly examine differences in biomarker expression across three subject groups: the NC group, the MCI group, and the AD group. This study enables us to assess the biomarker’s relevance across a broader spectrum of cognitive health, from normal aging through mild impairment to confirmed Alzheimer’s Disease, allowing for a finer understanding of its potential as a diagnostic indicator.

Furthermore, using the HCP dataset, we conduct two additional analyses focused on demographic and behavioral factors. The first is a sex difference analysis, where we compare biomarker expression between male and female participants. The second analysis examines differences in sleep quality, utilizing the Pittsburgh Sleep Quality Index (PSQI), which evaluates various dimensions of sleep, including duration, latency, efficiency, and disturbances. The PSQI scores can be ranked into three levels: scores between 0 ≤ PSQI ≤ 5 indicate good sleep quality; scores between 6 ≤ PSQI ≤ 10 reflect moderate sleep quality; and scores of PSQI ≥ 11 represent bad sleep quality. This study allows us to evaluate the biomarker’s sensitivity to sleep-related factors, which are increasingly acknowledged for their significant impact on cognitive health and brain function.

#### 3.2.2 Experimental Results

The *p*-values from the two-tailed Mann-Whitney U tests for NC vs. EMCI and EMCI vs. LMCI on the ADNI dataset are presented in Table 1. The results show that the *p*-values for NC vs. EMCI and EMCI vs. LMCI are both less than 0.05, indicating significant differences between these subject groups. These findings suggest that our proposed functional biomarker, instantaneous frequency, effectively distinguishes among NC, EMCI, and LMCI groups. Similarly, Table 2 presents the two-tailed *p*-values for NC vs. MCI and MCI vs. AD on the OASIS dataset. Additionally, Table 3 displays the two-tailed *p*-values for Male vs. Female, Good vs. Moderate sleep quality, and Moderate vs. Poor sleep quality on the HCP dataset. In all cases, the *p*-values are less than 0.05, clearly demonstrating that our proposed functional biomarkers reveal significant differences between these subgroups.

**Table 1:** The two-tailed *p*-values of EMCI vs NC and LMCI vs EMCI on ADNI dataset.

|  | NC | EMCI | LMCI |
| --- | --- | --- | --- |
| **EMCI-NC** | 1.0898e-47 | 1.0926e-24 | - |
| **LMCI-EMCI** | - | 4.6464e-07 | 6.2183e-04 |

**Table 2:** The two-tailed *p*-values of MCI vs NC and AD vs MCI on OASIS data.

|  | NC | MCI | AD |
| --- | --- | --- | --- |
| **MCI-NC** | 1.2743e-96 | 1.6747e-241 | - |
| **AD-MCI** | - | 7.7449e-12 | 6.7276e-05 |

**Table 3:**
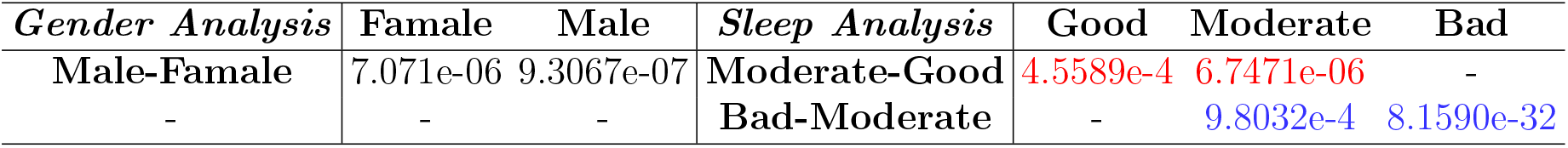
The two-tailed *p*-values of gender group analysis and sleep quality analysis on HCP dataset.

| *Gender Analysis* | Famale | Male | *Sleep Analysis* | Good | Moderate | Bad |
| --- | --- | --- | --- | --- | --- | --- |
| **Male-Famale** | 7.071e-06 | 9.3067e-07 | **Moderate-Good** | 4.5589e-4 | 6.7471e-06 | - |
| - | - | - | **Bad-Moderate** | - | 9.8032e-4 | 8.1590e-32 |

### 3.3 Identification of Distinct Connectomes Across Groups

We perform a connectome-level analysis to identify the significant brain connectomes differentiated by our proposed functional biomarkers when comparing different subject groups. Specifically, we use two-tailed Mann-Whitney U tests to evaluate differences in instantaneous frequency sequences for each brain connectome across subject groups. A connectome is considered distinct between two groups if its instantaneous frequency shows a significant difference. To control for multiple comparisons, the two-sided *p*-values are compared against the threshold 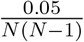, where *N* (*N* − 1) represents the total number of brain connectomes.

The group-specific distinct brain connectomes identified by the connectome instantaneous frequency are visualized in Figure 2 for the ADNI dataset, Figure 3 for the OASIS dataset, and Figure 4 for the HCP dataset. In the ADNI analysis (Figure 2), 680 brain connectomes show significant differences between the EMCI and LMCI groups, while 756 connectomes differ significantly between the NC and EMCI groups. The specific connectomes are annotated in Figure 2, with the corresponding brain ROI names provided in the Supplementary Materials.

**Figure 1:**
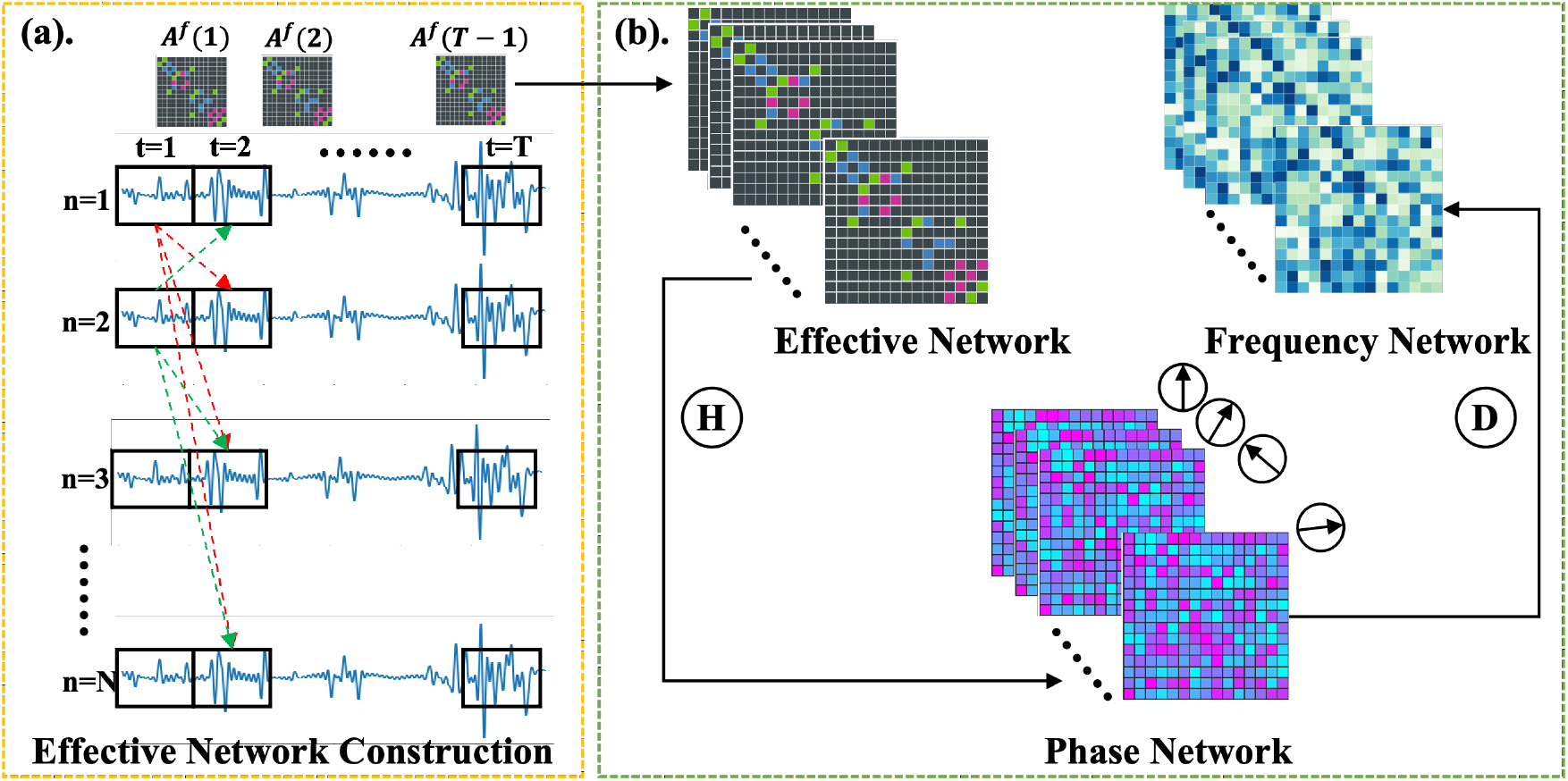
Diagram of the proposed method for computing the instantaneous frequency (IF) of brain connectomes, including **(a)** effective network construction and **(b)** IF network construction.

**Figure 2:**
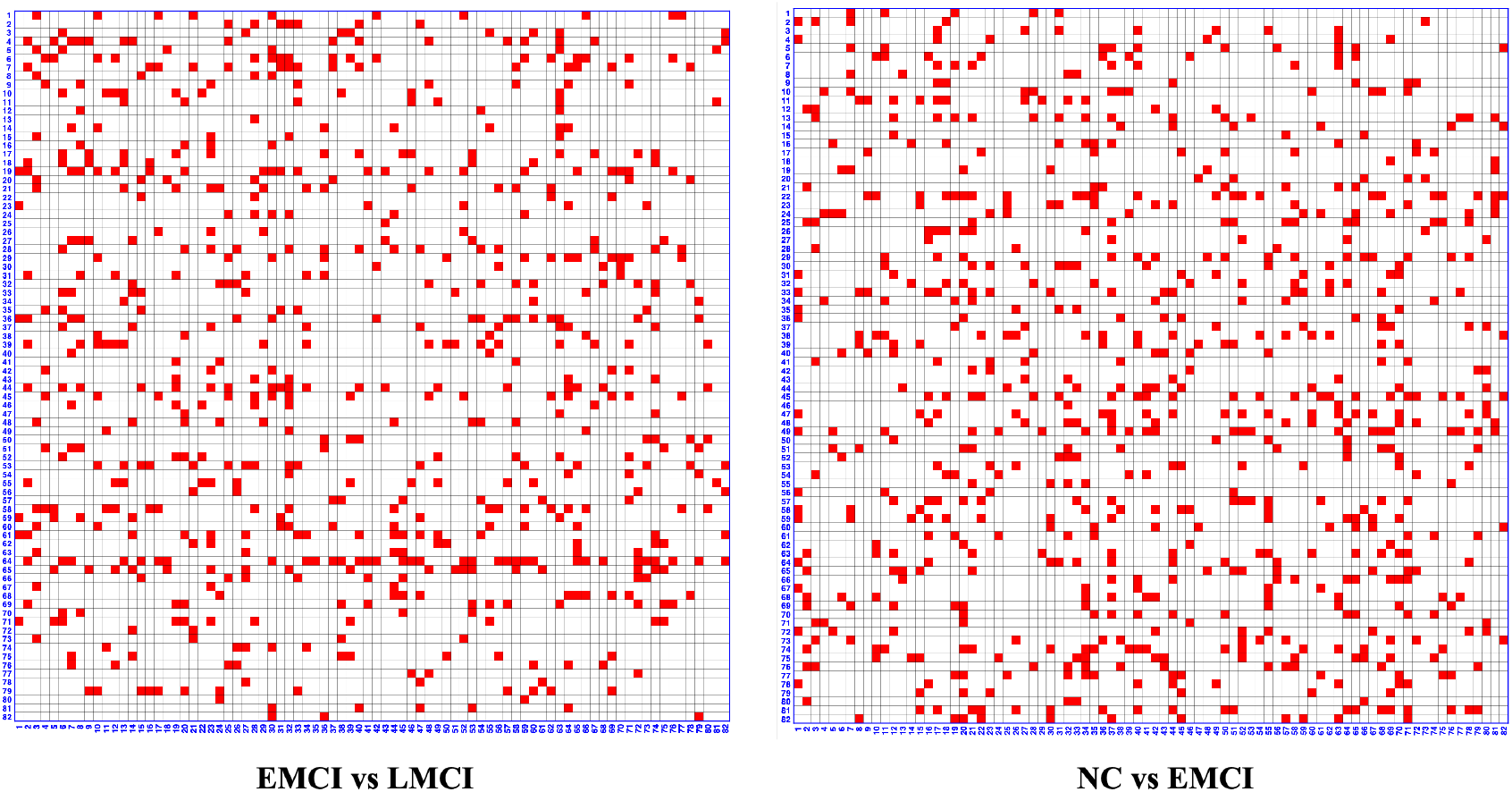
Significantly brain connectomes identified to distinguish EMCI and LMCI, and to distinguish NC and EMCI on ADNI dataset.

**Figure 3:**
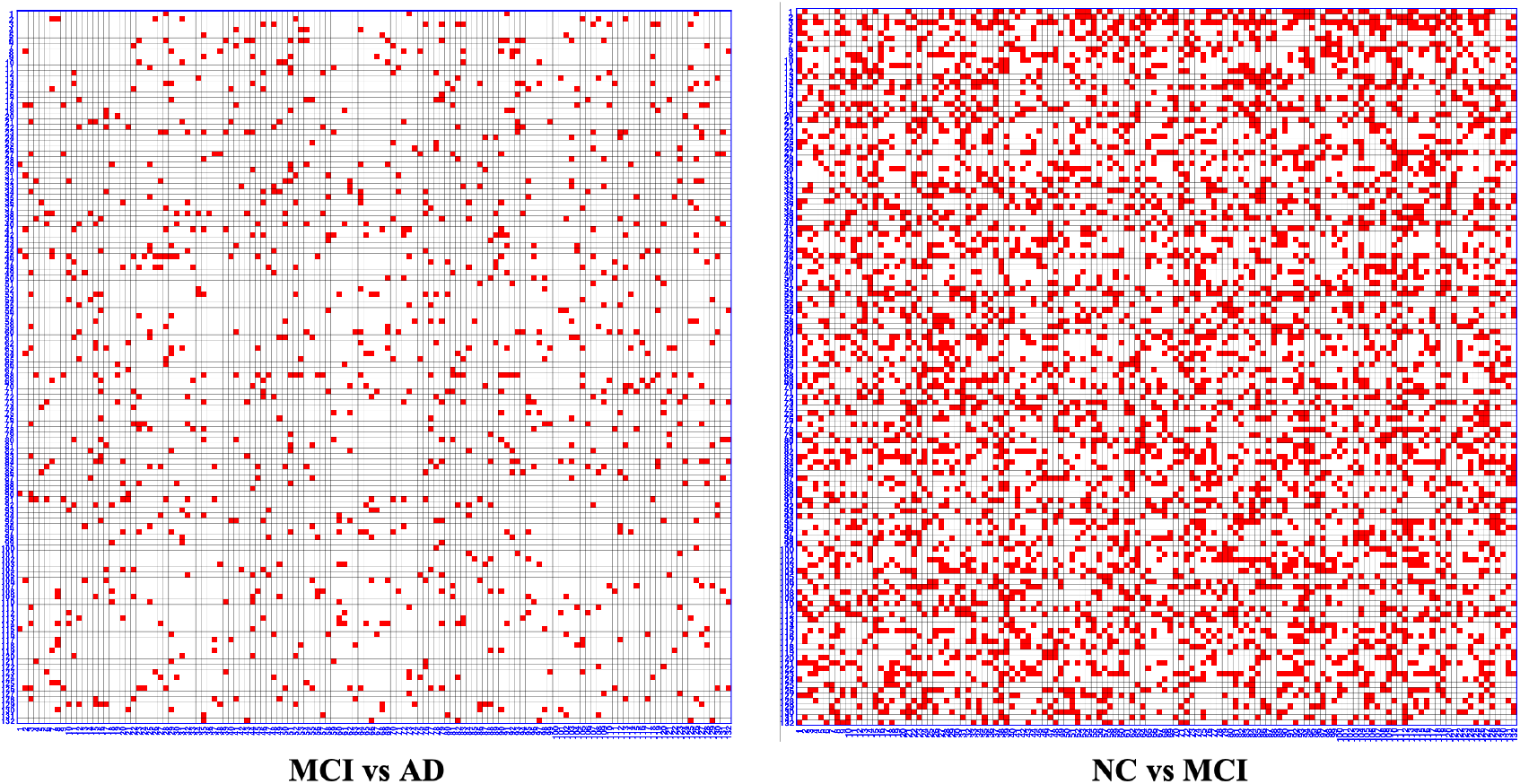
Significantly brain connectomes identified to distinguish MCI and AD, and to distinguish NC and MCI on OASIS dataset.

**Figure 4:**
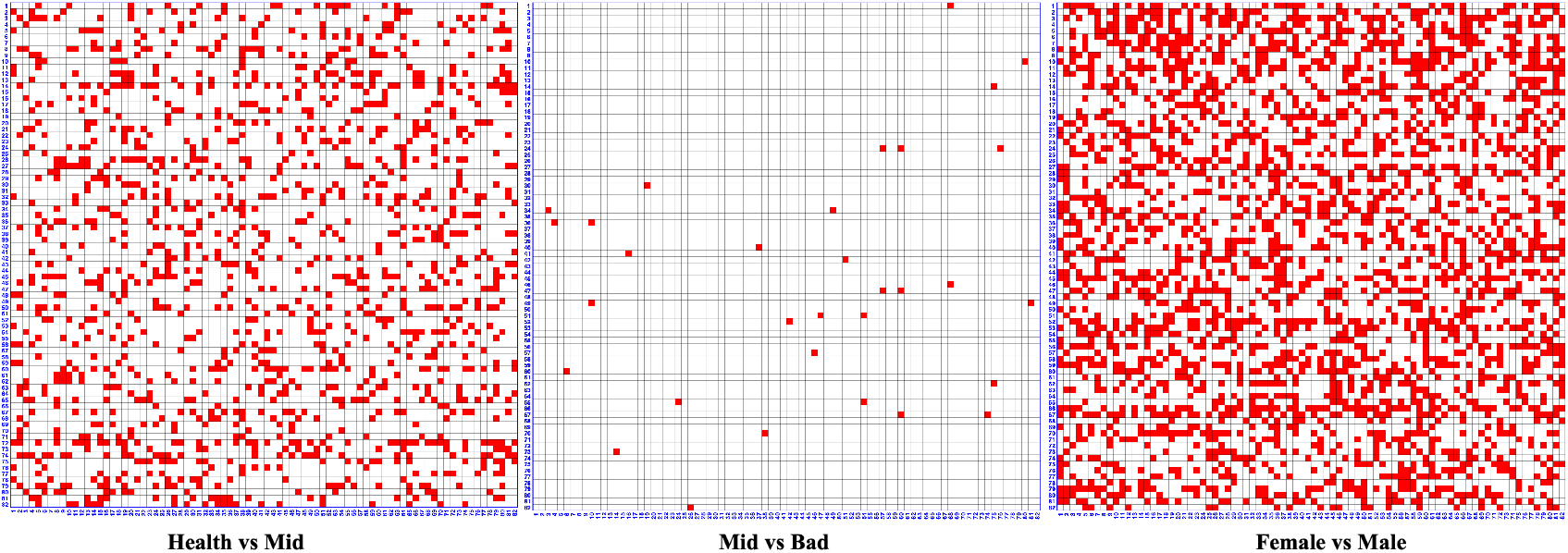
Significantly brain connectomes identified to distinguish Female and Male, and to distinguish different sleep quality states on HCP dataset.

For the OASIS dataset (see Figure 3), 974 brain connectomes show significant differences between the MCI and AD groups, and 4865 connectomes are significantly different between the NC and MCI groups. Similarly, the corresponding connectome details are located in Figure 3 and the related brain ROI names are included in the Supplementary Materials.

In the HCP dataset analysis (see Figure 4), 2495 connectomes are significantly different between male and female groups, 1296 connectomes differ significantly between good and moderate sleep quality groups, and only 32 connectomes differ significantly between moderate and poor sleep quality groups. Detailed connectome locations and corresponding ROI names are provided in the Supplementary Materials.

### 3.4 Visualizations of Identified Connectomes and Discussions

We compute the mean instantaneous frequency (i.e.,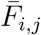) of the identified connectomes over the group subjects as follow:

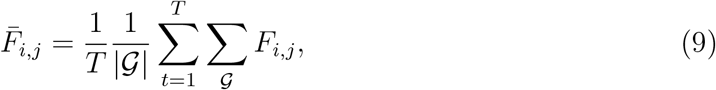

where *T* is the number of instantaneous frequency point and |𝒢| is the number of subjects in each group. Then we visualize the mean instantaneous frequency of the identified connectomes for the EMCI vs LMCI and for the NC vs EMCI on ADNI dataset in Figure 5. Similarly, the instantaneous frequency differences for the MCI vs AD, and NC vs MCI on OASIS dataset are shown in Figure 6. The instantaneous frequency differences for the male vs female, and different sleep qualities are shown in Figure 7.

**Figure 5:**
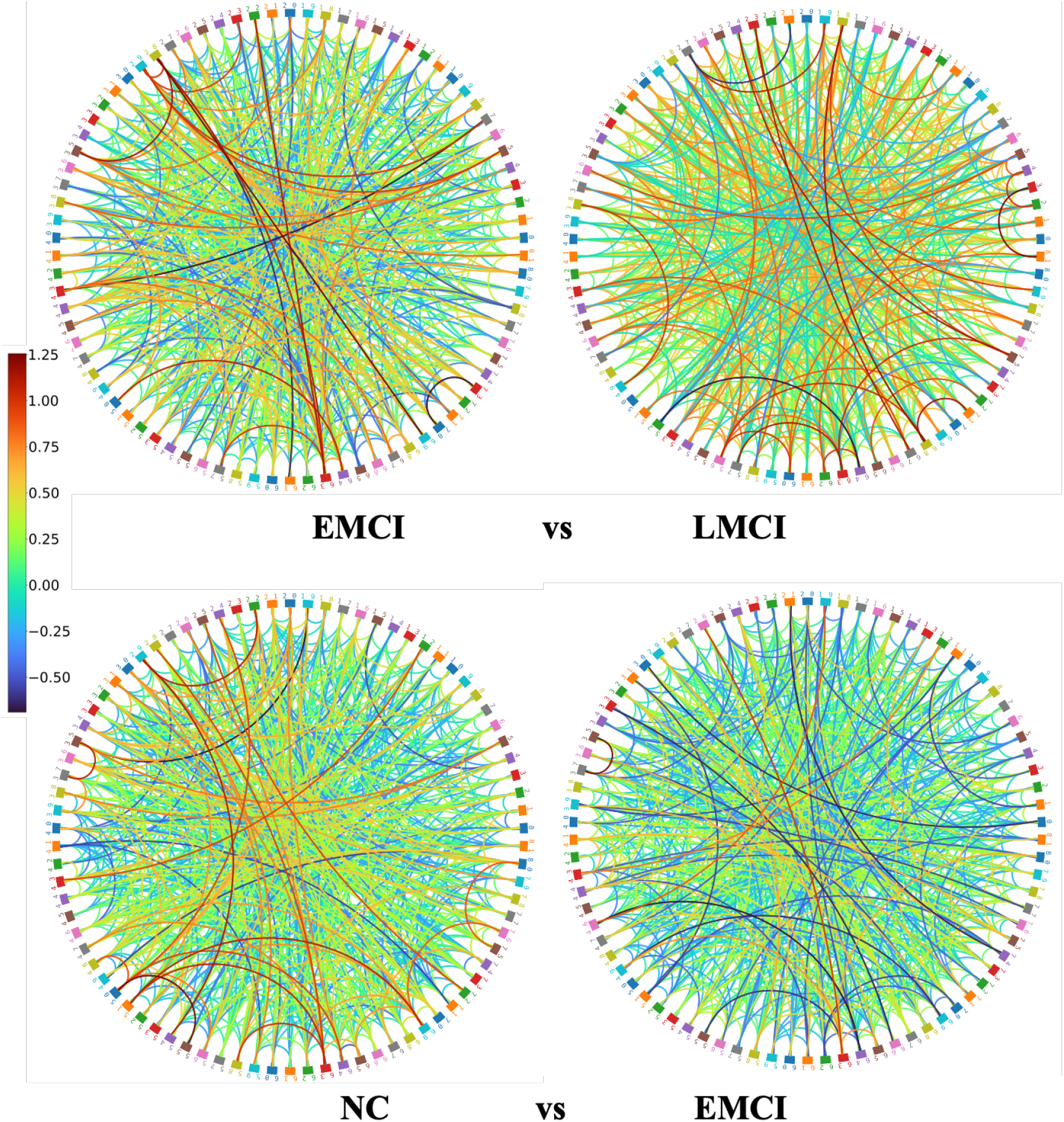
Mean instantaneous frequency 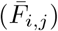 of the identified connectomes for EMCI vs LMCI and NC vs EMCI on ADNI dataset.

**Figure 6:**
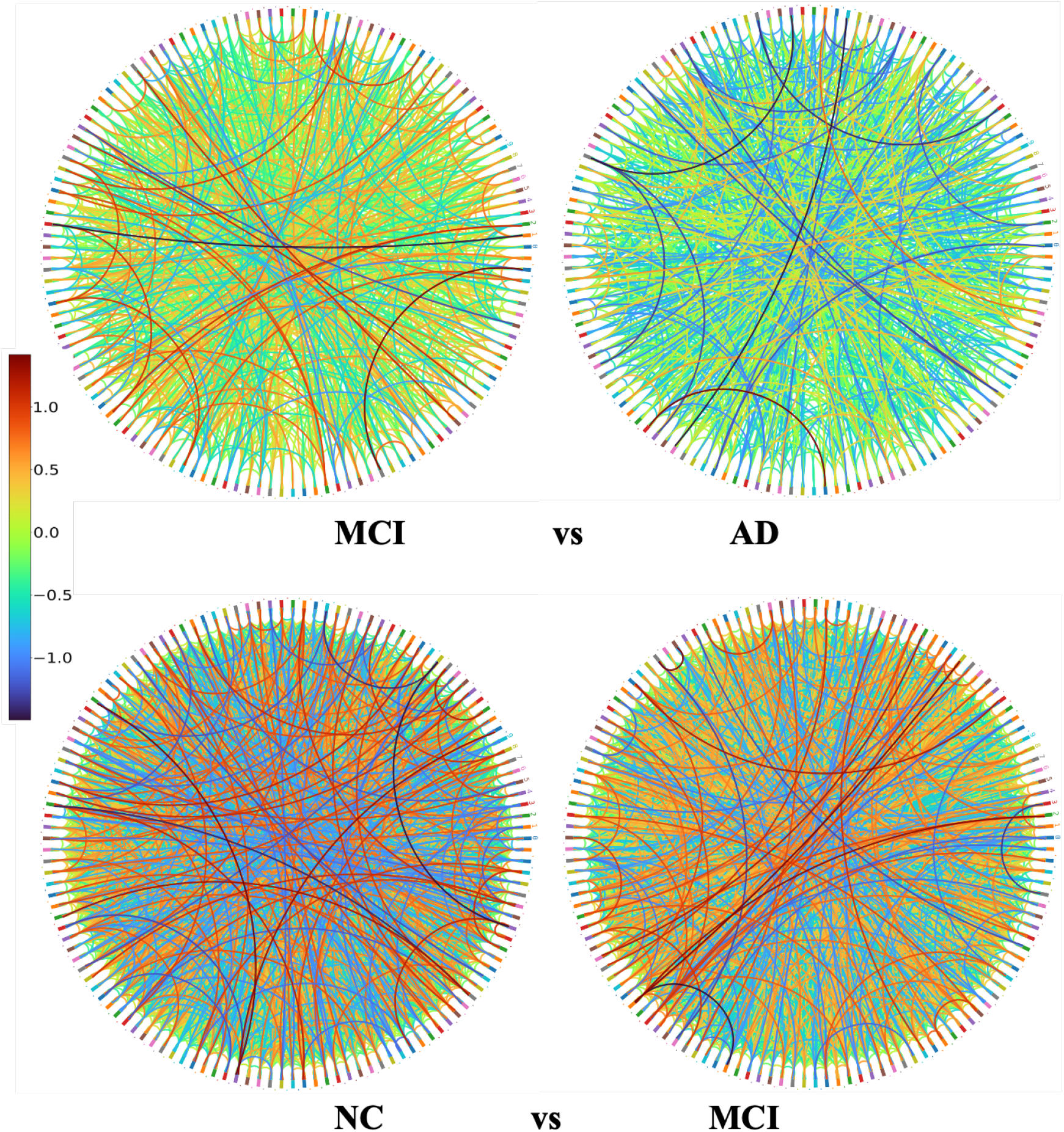
Mean instantaneous frequency 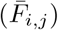 of the identified connectomes for MCI vs AD and NC vs MCI on OASIS dataset.

**Figure 7:**
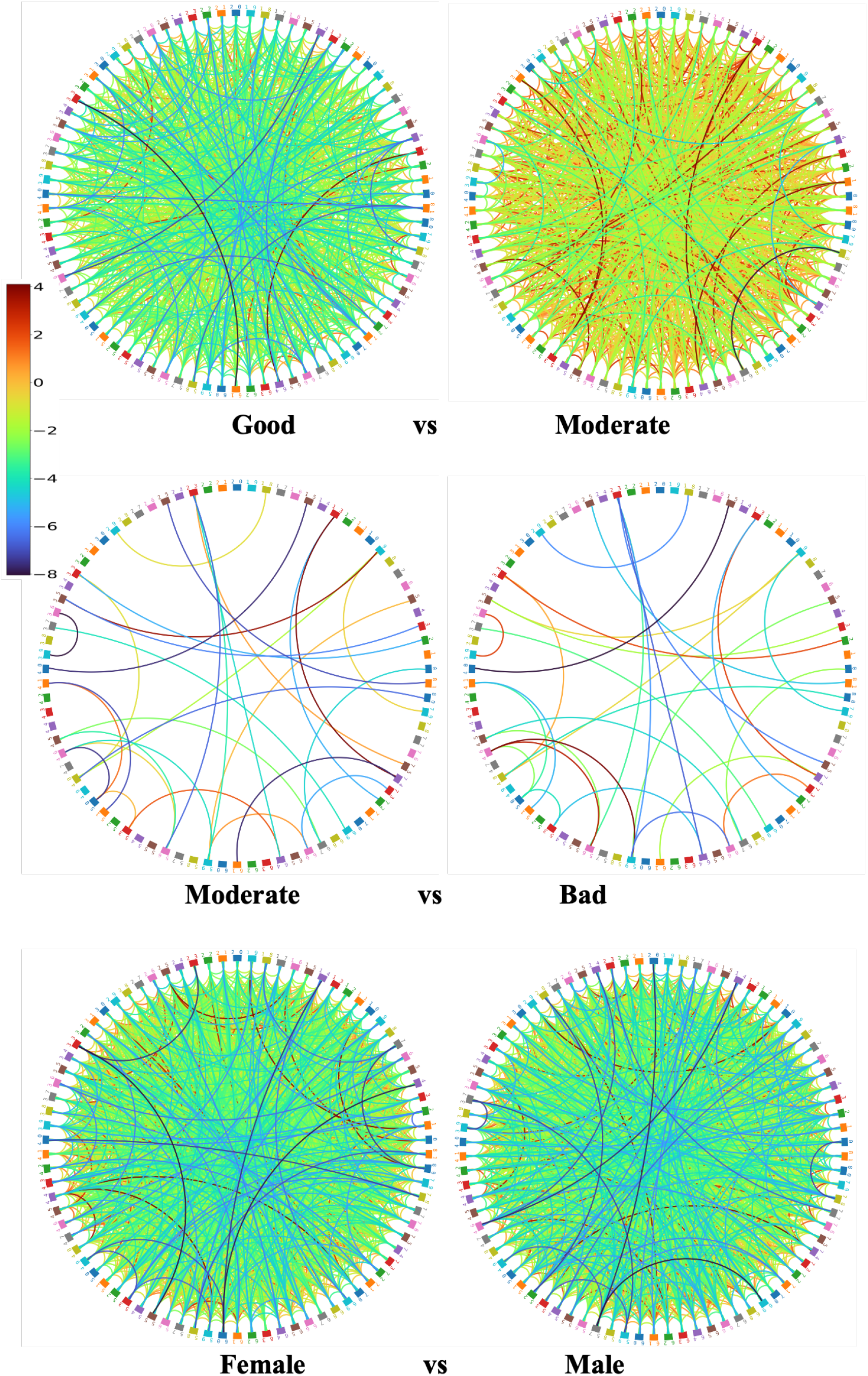
Mean instantaneous frequency 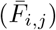 of the identified connectomes for Good vs Moderate, Moderate vs Bad sleep quality, and Female vs Male on HCP dataset.

Meanwhile, we present the mean and standard deviations of these 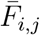 over the highlighted connectomes in the Figure 8. It shows in Figure 8 (a) and (b) that both the mean and variance of instantaneous frequency (IF) increase from NC to EMCI stage, and then decrease from EMCI to LMCI stage. This pattern reflects the brain’s compensatory mechanisms in early disease stages, followed by a decline in neural activity as the disease progresses. Previous studies have shown that during the early stages of AD, there is an increase in neural activity, particularly in the frontal regions. This hyperactivity is thought to be a compensatory response to the initial neuronal damage, aiming to maintain cognitive function [28, 29]. As the cognitive impairment progresses, the initial compensatory increase in neural activity is disrupted, resulting in a decline in neural activity compared to healthy aging. The brain’s reorganization observed in early stages is no longer sufficient to counteract the advancing pathology [30]. Figure 8 (c) shows that poor sleep quality results in an increase of the mean and variance of the IF, indicating greater instability in neural oscillations. This aligns with the fragmented sleep architecture observed in poor sleepers, reflecting inconsistent neural firing patterns, contributing to the subjective experience of poor sleep quality [31, 32]. In Figure 8, it shows that the male’s brain is more active than the females one, while the female’s brain is more stable.

**Figure 8:**
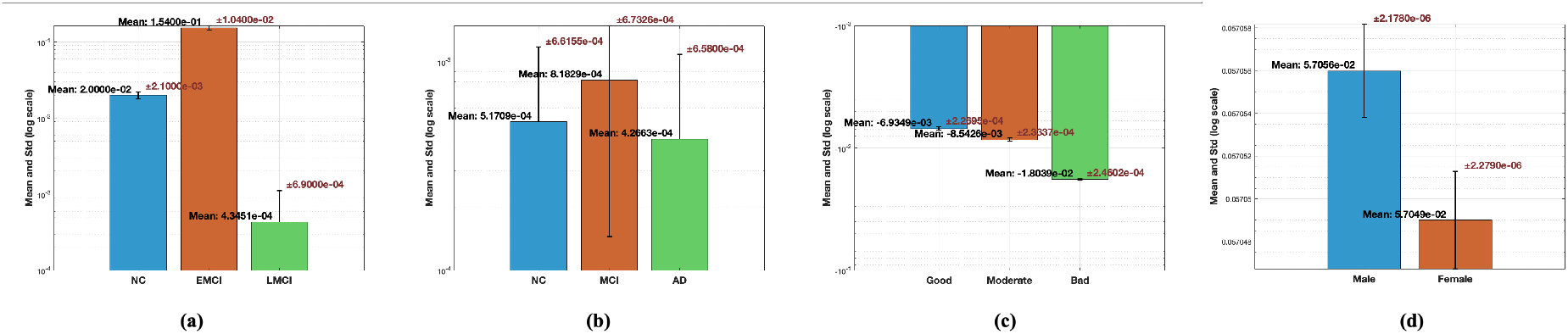
Mean and *s.t.d* values of 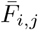 for the cognitive disease analyses on **(a)** ADNI dataset and **(b)** OASIS dataset, as well as for gender and sleep quality analyses on **(c)** HCP dataset.

## 4. Conclusion

This study introduces instantaneous frequency (IF) as a novel functional biomarker derived from dynamic brain effective connectivity. IF provides a robust measure of temporal fluctuations in brain network dynamics, offering insights into the mechanisms underlying cognitive decline, sex differences, and variations in sleep quality. Validation using ADNI, OASIS, and HCP datasets demonstrates IF’s ability to significantly discriminate between clinical and demographic groups. These findings suggest IF’s potential utility in exploring compensatory neural mechanisms, identifying disease-specific neural circuitry, and characterizing individual variability in brain dynamics. Future research should explore the integration of this biomarker into clinical diagnostic pipelines and its application to other neurological and psychiatric conditions.

## Fundings

This study was partially supported by the NIH (R01MH125928, and U01AG0 68057), the NSF (IIS 2319450 and IIS 2045848), as well as by the Presidential Research Fellowship (PRF) in the Department of Computer Science at the University of Texas Rio Grande Valley (UTRGV), and the UTRGV seed grant.

## Computation Resources

Part of the work utilized Bridges-2 [33] at the Pittsburgh Supercomputing Center through the ACCESS program, supported by NSF grants #2138259, #2138286, #2138307, #2137603, and #2138296. We also acknowledge the High Performance Computing Resource at the UTRGV. This cluster is based upon work supported by the National Science Foundation, MRI: Acquisition of a GPU-Accelerated Cluster, High Performance Rio Grande Valley Cluster (HiRGV), grant number 2018900, as well as the Department of Defense, HBC/MI under grant number W911NF2110169, and NSF IIS-2334389. “CAP: STARTER: South Texas AI Research, Training, and Education Resource.”

## ADNI Dataset

Data used in the preparation of this article were obtained from the Alzheimer’s Disease Neuroimaging Initiative (ADNI) database (adni.loni.usc.edu). The ADNI was launched in 2003 as a public-private partnership, led by Principal Investigator Michael W. Weiner, MD. The original goal of ADNI was to test whether serial magnetic resonance imaging (MRI), positron emission tomography (PET), other biological markers, and clinical and neuropsychological assessment can be combined to measure the progression of mild cognitive impairment (MCI) and early Alzheimer’s disease (AD). The current goals include validating biomarkers for clinical trials, improving the generalizability of ADNI data by increasing diversity in the participant cohort, and to provide data concerning the diagnosis and progression of Alzheimer’s disease to the scientific community. For up-to-date information, see adni.loni.usc.edu.

Data collection and sharing for the Alzheimer’s Disease Neuroimaging Initiative (ADNI) is funded by the National Institute on Aging (National Institutes of Health Grant U19AG02 4904). The grantee organization is the Northern California Institute for Research and Education. In the past, ADNI has also received funding from the National Institute of Biomedical Imaging and Bioengineering, the Canadian Institutes of Health Research, and private sector contributions through the Foundation for the National Institutes of Health (FNIH) including generous contributions from the following: AbbVie, Alzheimer’s Association; Alzheimer’s Drug Discovery Foundation; Araclon Biotech; BioClinica, Inc.; Biogen; Bristol-Myers Squibb Company; CereSpir, Inc.; Cogstate; Eisai Inc.; Elan Pharmaceuticals, Inc.; Eli Lilly and Company; EuroImmun; F. Hoffmann-La Roche Ltd and its affiliated company Genentech, Inc.; Fujirebio; GE Healthcare; IXICO Ltd.; Janssen Alzheimer Immunotherapy Research & Development, LLC.; Johnson & Johnson Pharmaceutical Research & Development LLC.; Lumosity; Lundbeck; Merck & Co., Inc.; Meso Scale Diagnostics, LLC.; NeuroRx Research; Neurotrack Technologies; Novartis Pharmaceuticals Corporation; Pfizer Inc.; Piramal Imaging; Servier; Takeda Pharmaceutical Company; and Transition Therapeutics.

## HCP Dataset

Data collection and sharing for this project was provided by the Human Connectome Project (HCP; Principal Investigators: Bruce Rosen, M.D., Ph.D., Arthur W. Toga, Ph.D., Van J. Weeden, MD). HCP funding was provided by the National Institute of Dental and Craniofacial Research (NIDCR), the National Institute of Mental Health (NIMH), and the National Institute of Neurological Disorders and Stroke (NINDS). HCP data are disseminated by the Laboratory of Neuro Imaging at the University of Southern California

## OASIS Dataset

For OASIS, the magnetic resonance imaging and neuropsychological test data that support the findings of this study are available in [“OASIS-3”] (https://doi.org/10.1101/2019.12.13.19014902). OASIS-3: Longitudinal Multimodal Neuroimaging: Principal Investigators: T. Benzinger, D. Marcus, J. Morris; NIH grants P30 AG066444, P50 AG00561, P30 NS09857781, P01 AG026276, P01 AG030991, R01 AG043434, UL1 TR000448, R01 EB009352. AV-45 doses were provided by Avid Radiopharmaceuticals, a wholly owned subsidiary of Eli Lilly.

OASIS-3 data^1^ are openly available to the scientific community. Prior to accessing the data, users are required to agree to the OASIS Data Use Terms (DUT), which follow the Creative Commons Attribution 4.0 License.

## Supplementary Materials

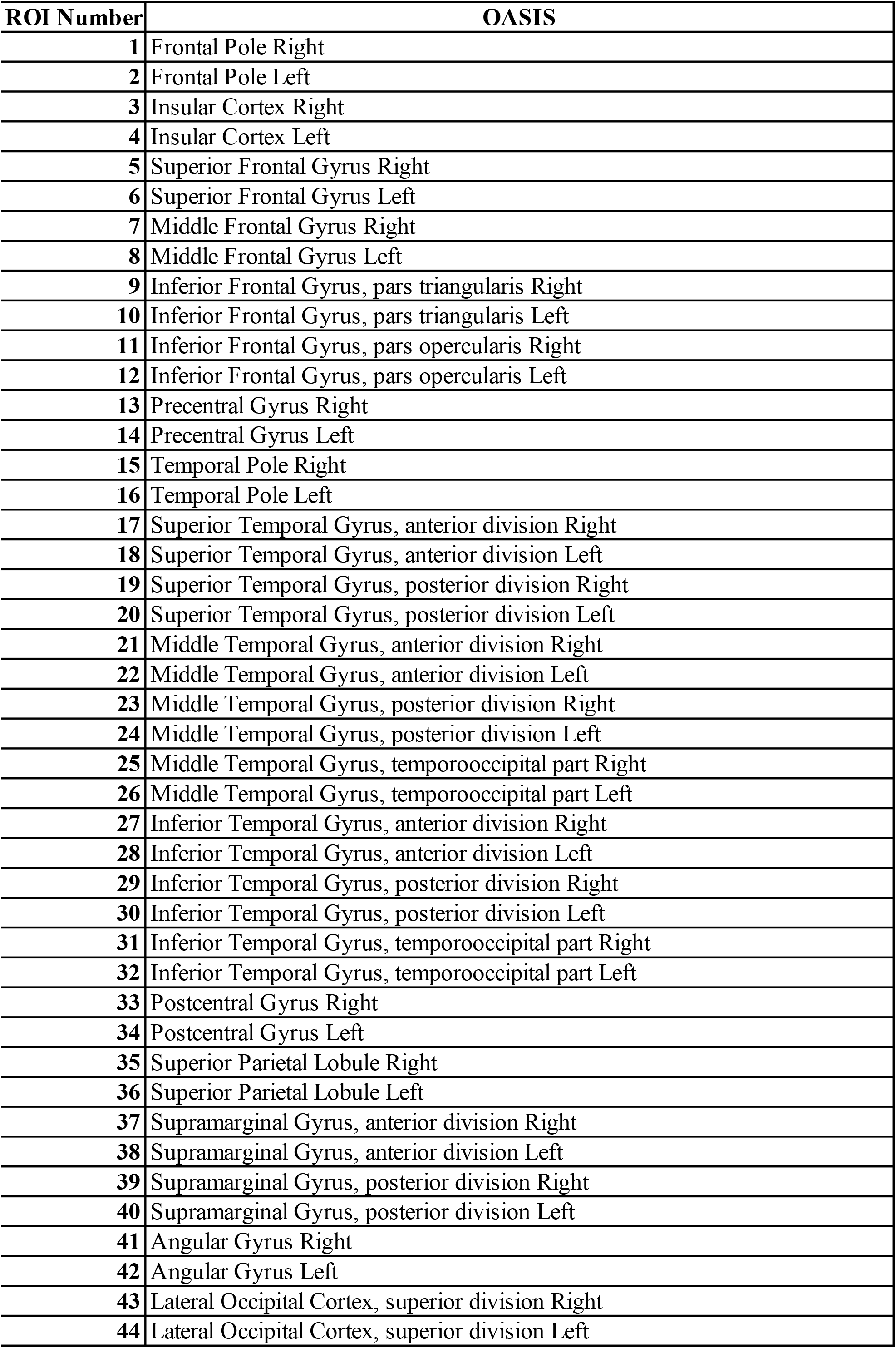
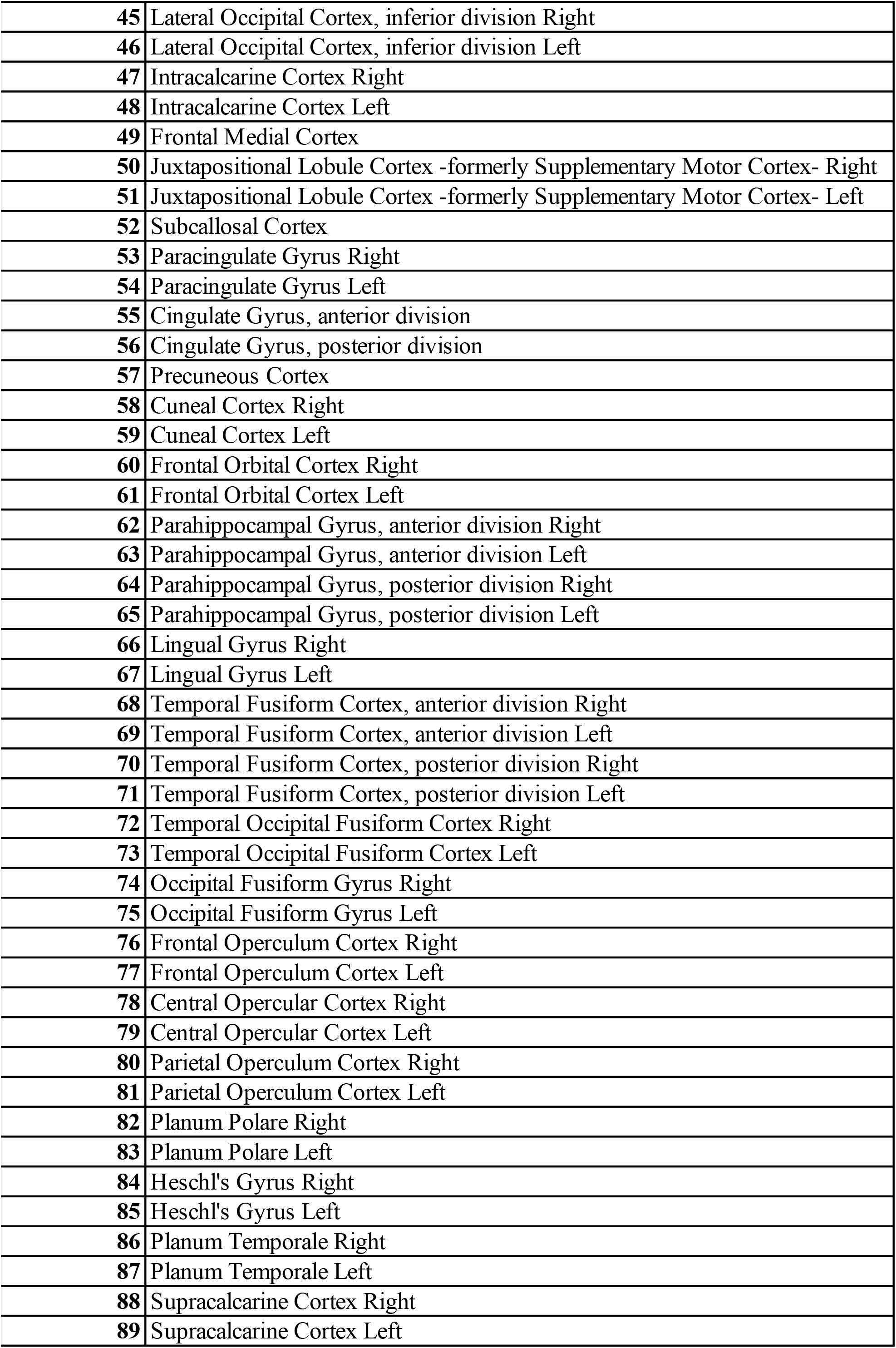
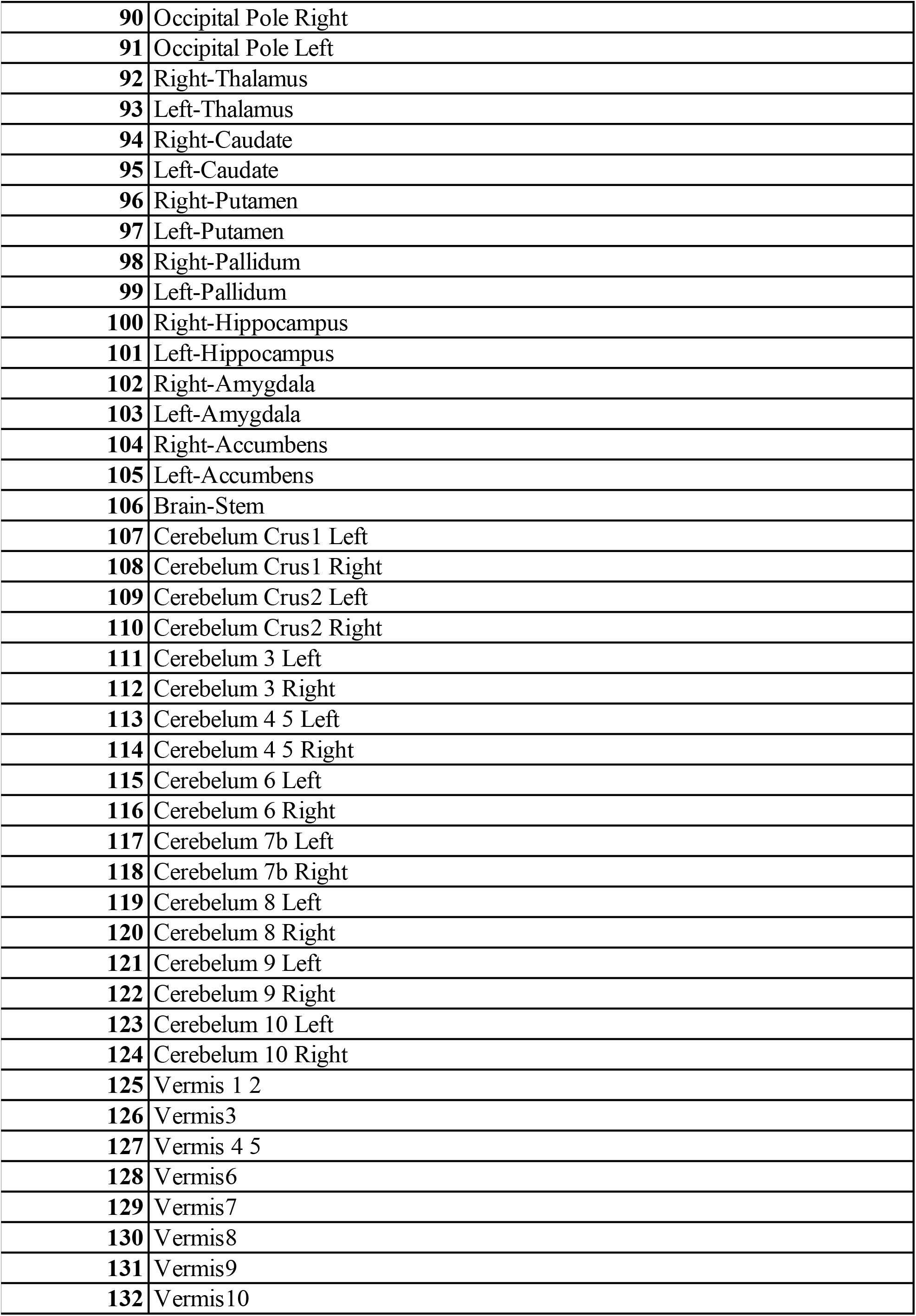

| ROI Number | OASIS |
| --- | --- |
| 1 | Frontal Pole Right |
| 2 | Frontal Pole Left |
| 3 | Insular Cortex Right |
| 4 | Insular Cortex Left |
| 5 | Superior Frontal Gyrus Right |
| 6 | Superior Frontal Gyrus Left |
| 7 | Middle Frontal Gyrus Right |
| 8 | Middle Frontal Gyrus Left |
| 9 | Inferior Frontal Gyrus, pars triangularis Right |
| 10 | Inferior Frontal Gyrus, pars triangularis Left |
| 11 | Inferior Frontal Gyrus, pars opercularis Right |
| 12 | Inferior Frontal Gyrus, pars opercularis Left |
| 13 | Precentral Gyrus Right |
| 14 | Precentral Gyrus Left |
| 15 | Temporal Pole Right |
| 16 | Temporal Pole Left |
| 17 | Superior Temporal Gyrus, anterior division Right |
| 18 | Superior Temporal Gyrus, anterior division Left |
| 19 | Superior Temporal Gyrus, posterior division Right |
| 20 | Superior Temporal Gyrus, posterior division Left |
| 21 | Middle Temporal Gyrus, anterior division Right |
| 22 | Middle Temporal Gyrus, anterior division Left |
| 23 | Middle Temporal Gyrus, posterior division Right |
| 24 | Middle Temporal Gyrus, posterior division Left |
| 25 | Middle Temporal Gyrus, temporooccipital part Right |
| 26 | Middle Temporal Gyrus, temporooccipital part Left |
| 27 | Inferior Temporal Gyrus, anterior division Right |
| 28 | Inferior Temporal Gyrus, anterior division Left |
| 29 | Inferior Temporal Gyrus, posterior division Right |
| 30 | Inferior Temporal Gyrus, posterior division Left |
| 31 | Inferior Temporal Gyrus, temporooccipital part Right |
| 32 | Inferior Temporal Gyrus, temporooccipital part Left |
| 33 | Postcentral Gyrus Right |
| 34 | Postcentral Gyrus Left |
| 35 | Superior Parietal Lobule Right |
| 36 | Superior Parietal Lobule Left |
| 37 | Supramarginal Gyrus, anterior division Right |
| 38 | Supramarginal Gyrus, anterior division Left |
| 39 | Supramarginal Gyrus, posterior division Right |
| 40 | Supramarginal Gyrus, posterior division Left |
| 41 | Angular Gyrus Right |
| 42 | Angular Gyrus Left |
| 43 | Lateral Occipital Cortex, superior division Right |
| 44 | Lateral Occipital Cortex, superior division Left |
| 45 | Lateral Occipital Cortex, inferior division Right |
| 46 | Lateral Occipital Cortex, inferior division Left |
| 47 | Intracalcarine Cortex Right |
| 48 | Intracalcarine Cortex Left |
| 49 | Frontal Medial Cortex |
| 50 | Juxtapositional Lobule Cortex -formerly Supplementary Motor Cortex- Right |
| 51 | Juxtapositional Lobule Cortex -formerly Supplementary Motor Cortex- Left |
| 52 | Subcallosal Cortex |
| 53 | Paracingulate Gyrus Right |
| 54 | Paracingulate Gyrus Left |
| 55 | Cingulate Gyrus, anterior division |
| 56 | Cingulate Gyrus, posterior division |
| 57 | Precuneous Cortex |
| 58 | Cuneal Cortex Right |
| 59 | Cuneal Cortex Left |
| 60 | Frontal Orbital Cortex Right |
| 61 | Frontal Orbital Cortex Left |
| 62 | Parahippocampal Gyrus, anterior division Right |
| 63 | Parahippocampal Gyrus, anterior division Left |
| 64 | Parahippocampal Gyrus, posterior division Right |
| 65 | Parahippocampal Gyrus, posterior division Left |
| 66 | Lingual Gyrus Right |
| 67 | Lingual Gyrus Left |
| 68 | Temporal Fusiform Cortex, anterior division Right |
| 69 | Temporal Fusiform Cortex, anterior division Left |
| 70 | Temporal Fusiform Cortex, posterior division Right |
| 71 | Temporal Fusiform Cortex, posterior division Left |
| 72 | Temporal Occipital Fusiform Cortex Right |
| 73 | Temporal Occipital Fusiform Cortex Left |
| 74 | Occipital Fusiform Gyrus Right |
| 75 | Occipital Fusiform Gyrus Left |
| 76 | Frontal Operculum Cortex Right |
| 77 | Frontal Operculum Cortex Left |
| 78 | Central Opercular Cortex Right |
| 79 | Central Opercular Cortex Left |
| 80 | Parietal Operculum Cortex Right |
| 81 | Parietal Operculum Cortex Left |
| 82 | Planum Polare Right |
| 83 | Planum Polare Left |
| 84 | Heschl's Gyrus Right |
| 85 | Heschl's Gyrus Left |
| 86 | Planum Temporale Right |
| 87 | Planum Temporale Left |
| 88 | Supracalcarine Cortex Right |
| 89 | Supracalcarine Cortex Left |
| **90** | Occipital Pole Right |
| **91** | Occipital Pole Left |
| **92** | Right-Thalamus |
| **93** | Left-Thalamus |
| **94** | Right-Caudate |
| **95** | Left-Caudate |
| **96** | Right-Putamen |
| **97** | Left-Putamen |
| **98** | Right-Pallidum |
| **99** | Left-Pallidum |
| **100** | Right-Hippocampus |
| **101** | Left-Hippocampus |
| **102** | Right-Amygdala |
| **103** | Left-Amygdala |
| **104** | Right-Accumbens |
| **105** | Left-Accumbens |
| **106** | Brain-Stem |
| **107** | Cerebelum Crus1 Left |
| **108** | Cerebelum Crus1 Right |
| **109** | Cerebelum Crus2 Left |
| **110** | Cerebelum Crus2 Right |
| **111** | Cerebelum 3 Left |
| **112** | Cerebelum 3 Right |
| **113** | Cerebelum 4 5 Left |
| **114** | Cerebelum 4 5 Right |
| **115** | Cerebelum 6 Left |
| **116** | Cerebelum 6 Right |
| **117** | Cerebelum 7b Left |
| **118** | Cerebelum 7b Right |
| **119** | Cerebelum 8 Left |
| **120** | Cerebelum 8 Right |
| **121** | Cerebelum 9 Left |
| **122** | Cerebelum 9 Right |
| **123** | Cerebelum 10 Left |
| **124** | Cerebelum 10 Right |
| **125** | Vermis 1 2 |
| **126** | Vermis3 |
| **127** | Vermis 4 5 |
| **128** | Vermis6 |
| **129** | Vermis7 |
| **130** | Vermis8 |
| **131** | Vermis9 |
| **132** | Vermis10 |

| ROI Number | HCP and ADNI |
| --- | --- |
| 1 | Left-Thalamus |
| 2 | Left-Caudate |
| 3 | Left-Putamen |
| 4 | Left-Pallidum |
| 5 | Left-Hippocampus |
| 6 | Left-Amygdala |
| 7 | Left-Accumbens-area |
| 8 | Right-Thalamus |
| 9 | Right-Caudate |
| 10 | Right-Putamen |
| 11 | Right-Pallidum |
| 12 | Right-Hippocampus |
| 13 | Right-Amygdala |
| 14 | Right-Accumbens-area |
| 15 | ctx-lh-bankssts |
| 16 | ctx-lh-caudalanteriorcingulate |
| 17 | ctx-lh-caudalmiddlefrontal |
| 18 | ctx-lh-cuneus |
| 19 | ctx-lh-entorhinal |
| 20 | ctx-lh-fusiform |
| 21 | ctx-lh-inferiorparietal |
| 22 | ctx-lh-inferiortemporal |
| 23 | ctx-lh-isthmuscingulate |
| 24 | ctx-lh-lateraloccipital |
| 25 | ctx-lh-lateralorbitofrontal |
| 26 | ctx-lh-lingual |
| 27 | ctx-lh-medialorbitofrontal |
| 28 | ctx-lh-middletemporal |
| 29 | ctx-lh-parahippocampal |
| 30 | ctx-lh-paracentral |
| 31 | ctx-lh-parsopercularis |
| 32 | ctx-lh-parsorbitalis |
| 33 | ctx-lh-parstriangularis |
| 34 | ctx-lh-pericalcarine |
| 35 | ctx-lh-postcentral |
| 36 | ctx-lh-posteriorcingulate |
| 37 | ctx-lh-precentral |
| 38 | ctx-lh-precuneus |
| 39 | ctx-lh-rostralanteriorcingulate |
| 40 | ctx-lh-rostralmiddlefrontal |
| 41 | ctx-lh-superiorfrontal |
| 42 | ctx-lh-superiorparietal |
| 43 | ctx-lh-superiortemporal |
| 44 | ctx-lh-supramarginal |
| 45 | ctx-lh-frontalpole |
| 46 | ctx-lh-temporalpole |
| 47 | ctx-lh-transversetemporal |
| 48 | ctx-lh-insula |
| 49 | ctx-rh-bankssts |
| 50 | ctx-rh-caudalanteriorcingulate |
| 51 | ctx-rh-caudalmiddlefrontal |
| 52 | ctx-rh-cuneus |
| 53 | ctx-rh-entorhinal |
| 54 | ctx-rh-fusiform |
| 55 | ctx-rh-inferiorparietal |
| 56 | ctx-rh-inferiortemporal |
| 57 | ctx-rh-isthmuscingulate |
| 58 | ctx-rh-lateraloccipital |
| 59 | ctx-rh-lateralorbitofrontal |
| 60 | ctx-rh-lingual |
| 61 | ctx-rh-medialorbitofrontal |
| 62 | ctx-rh-middletemporal |
| 63 | ctx-rh-parahippocampal |
| 64 | ctx-rh-paracentral |
| 65 | ctx-rh-parsopercularis |
| 66 | ctx-rh-parsorbitalis |
| 67 | ctx-rh-parstriangularis |
| 68 | ctx-rh-pericalcarine |
| 69 | ctx-rh-postcentral |
| 70 | ctx-rh-posteriorcingulate |
| 71 | ctx-rh-precentral |
| 72 | ctx-rh-precuneus |
| 73 | ctx-rh-rostralanteriorcingulate |
| 74 | ctx-rh-rostralmiddlefrontal |
| 75 | ctx-rh-superiorfrontal |
| 76 | ctx-rh-superiorparietal |
| 77 | ctx-rh-superiortemporal |
| 78 | ctx-rh-supramarginal |
| 79 | ctx-rh-frontalpole |
| 80 | ctx-rh-temporalpole |
| 81 | ctx-rh-transversetemporal |
| 82 | ctx-rh-insula |

## Footnotes

1 https://www.oasisbrains.org

## Notes

### Competing Interest Statement

The authors have declared no competing interest.

## References

[1] C. R. Jack, D. S. Knopman, W. J. Jagust, L. M. Shaw, P. S. Aisen, M. W. Weiner, R. C. Petersen, J. Q. Trojanowski, Hypothetical model of dynamic biomarkers of the alzheimer’s pathological cascade, The Lancet Neurology 9 (2010) 119–128.

[2] H. Tang, L. Guo, X. Fu, B. Qu, P. M. Thompson, H. Huang, L. Zhan, Hierarchical brain embedding using explainable graph learning, in: 2022 IEEE 19th International Symposium on Biomedical Imaging (ISBI), IEEE, 2022, pp. 1–5.

[3] C. J. Honey, R. Kötter, M. Breakspear, O. Sporns, Network structure of cerebral cortex shapes functional connectivity on multiple time scales, Proceedings of the National Academy of Sciences 104 (2007) 10240–10245.

[4] D. S. Bassett, N. F. Wymbs, M. A. Porter, P. J. Mucha, J. M. Carlson, S. T. Grafton, Dynamic reconfiguration of human brain networks during learning, Proceedings of the National Academy of Sciences 108 (2011) 7641–7646.

[5] H. Tang, L. Guo, X. Fu, Y. Wang, S. Mackin, O. Ajilore, A. D. Leow, P. M. Thompson, H. Huang, L. Zhan, Signed graph representation learning for functional-to-structural brain network mapping, Medical image analysis 83 (2023) 102674.

[6] K. Ye, H. Tang, S. Dai, L. Guo, J. Y. Liu, Y. Wang, A. Leow, P. M. Thompson, H. Huang, L. Zhan, Bidirectional mapping with contrastive learning on multimodal neuroimaging data, in: H. Greenspan, A. Madabhushi, P. Mousavi, S. Salcudean, J. Duncan, T. Syeda-Mahmood, R. Taylor (Eds.), International Conference on Medical Image Computing and Computer-Assisted Intervention, Springer, 2023, pp. 138–148.

[7] K. J. Friston, Functional and effective connectivity: a review, Brain connectivity 1 (2011) 13–36.

[8] H. Tang, G. Ma, L. Guo, X. Fu, H. Huang, L. Zhan, Contrastive brain network learning via hierarchical signed graph pooling model, IEEE transactions on neural networks and learning systems (2022).

[9] H. Tang, G. Liu, S. Dai, K. Ye, K. Zhao, W. Wang, C. Yang, L. He, A. Leow, P. Thompson, et al., Interpretable spatio-temporal embedding for brain structural-effective network with ordinary differential equation, in: International Conference on Medical Image Computing and Computer-Assisted Intervention, Springer, 2024, pp. 227–237.

[10] A. Raj, A. Kuceyeski, M. Weiner, A network diffusion model of disease progression in dementia, Neuron 73 (2012) 1204–1215.

[11] C. R. Jack Jr, D. S. Knopman, W. J. Jagust, R. C. Petersen, M. W. Weiner, P. S. Aisen, L. M. Shaw, P. Vemuri, H. J. Wiste, S. D. Weigand, et al., Update on hypothetical model of alzheimer’s disease biomarkers, Lancet neurology 12 (2013) 207.

[12] O. Speck, T. Ernst, J. Braun, C. Koch, E. Miller, L. Chang, Gender differences in the functional organization of the brain for working memory, Neuroreport 11 (2000) 2581–2585.

[13] A. J. Krause, E. B. Simon, B. A. Mander, S. M. Greer, J. M. Saletin, A. N. Goldstein-Piekarski, M. P. Walker, The sleep-deprived human brain, Nature Reviews Neuroscience 18 (2017) 404–418.

[14] A. C. Marreiros, K. E. Stephan, K. J. Friston, Dynamic causal modeling, Scholarpedia 5 (2010) 9568.

[15] K. E. Stephan, W. D. Penny, R. J. Moran, H. E. den Ouden, J. Daunizeau, K. J. Friston, Ten simple rules for dynamic causal modeling, Neuroimage 49 (2010) 3099–3109.

[16] R. Sanchez-Romero, J. D. Ramsey, K. Zhang, M. R. Glymour, B. Huang, C. Glymour, Estimating feedforward and feedback effective connections from fmri time series: Assessments of statistical methods, Network Neuroscience 3 (2019) 274–306.

[17] S. M. Smith, K. L. Miller, G. Salimi-Khorshidi, M. Webster, C. F. Beckmann, T. E. Nichols, J. D. Ramsey, M. W. Woolrich, Network modelling methods for fmri, Neuroimage 54 (2011) 875–891.

[18] B. Boashash, Estimating and interpreting the instantaneous frequency of a signal. i. fundamentals, Proceedings of the IEEE 80 (1992) 520–538.

[19] L. Cohen, Time-frequency analysis, volume 778, Prentice Hall PTR New Jersey, 1995.

[20] D. Gabor, Theory of communication. part 1: The analysis of information, Journal of the Institution of Electrical Engineers-part III: radio and communication engineering 93 (1946) 429–441.

[21] H. Sakoe, S. Chiba, Dynamic programming algorithm optimization for spoken word recognition, IEEE transactions on acoustics, speech, and signal processing 26 (1978) 43–49.

[22] E. Keogh, C. A. Ratanamahatana, Exact indexing of dynamic time warping, Knowledge and information systems 7 (2005) 358–386.

[23] D. J. Berndt, J. Clifford, Using dynamic time warping to find patterns in time series, in: U. M. Fayyad, R. Uthurusamy (Eds.), Proceedings of the 3rd international conference on knowledge discovery and data mining, 1994, pp. 359–370.

[24] H. B. Mann, D. R. Whitney, On a test of whether one of two random variables is stochastically larger than the other, The annals of mathematical statistics (1947) 50–60.

[25] N. Nachar, et al., The mann-whitney u: A test for assessing whether two independent samples come from the same distribution, Tutorials in quantitative Methods for Psychology 4 (2008) 13–20.

[26] C. Misra, Y. Fan, C. Davatzikos, Baseline and longitudinal patterns of brain atrophy in mci patients, and their use in prediction of short-term conversion to ad: results from adni, Neuroimage 44 (2009) 1415–1422.

[27] D. C. Van Essen, S. M. Smith, D. M. Barch, T. E. Behrens, E. Yacoub, K. Ugurbil, W.-M. H. Consortium, et al., The wu-minn human connectome project: an overview, Neuroimage 80 (2013) 62–79.

[28] I. Aganj, A. Frau-Pascual, J. E. Iglesias, A. Yendiki, J. C. Augustinack, D. H. Salat, B. Fischl, Compensatory brain connection discovery in alzheimer’s disease, in: 2020 IEEE 17th International Symposium on Biomedical Imaging (ISBI), IEEE, 2020, pp. 283–287.

[29] S. Gaubert, F. Raimondo, M. Houot, M.-C. Corsi, L. Naccache, J. Diego Sitt, B. Hermann, D. Oudiette, G. Gagliardi, M.-O. Habert, et al., Eeg evidence of compensatory mechanisms in preclinical alzheimer’s disease, Brain 142 (2019) 2096–2112.

[30] J. Gallego-Rudolf, A. I. Wiesman, A. Pichet Binette, S. Villeneuve, S. Baillet, P.-A. research group, Synergistic association of aβ and tau pathology with cortical neurophysiology and cognitive decline in asymptomatic older adults, Nature Neuroscience (2024) 1–8.

[31] W. Cai, L. Chen, Y. Dai, B. Chen, D. Zheng, Y. Li, Association between eeg power during sleep and attention levels in patients with major depressive disorder, Nature and Science of Sleep (2024) 855–864.

[32] X.-J. Dai, J. Zhang, Y. Wang, Y. Ma, K. Shi, Eeg and fmri for sleep and sleep disorders–mechanisms and clinical implications, 2021.

[33] S. T. Brown, P. Buitrago, E. Hanna, S. Sanielevici, R. Scibek, N. A. Nystrom, Bridges-2: A platform for rapidly-evolving and data intensive research, in: Practice and Experience in Advanced Research Computing, 2021, pp. 1–4.

